# Effects of Full-cycle Exposure to Difenoconazole in Parental Zebrafish on the Liver-gut Axis of F0 and F1 Generations

**DOI:** 10.1101/2022.04.29.490113

**Authors:** Tao Zhu, Siwen Wang, Dong Li

## Abstract

To explore the effects of long-term exposure to low-dose difenoconazole (DCZ) on lipid metabolism in the liver-gut axis, we exposed zebrafish embryos to ambient concentrations of DCZ for 120 days and comprehensively analyzed the metabolic and microbial networks of the F0 and F1 generations using biochemical, metabolomic and metagenomics approaches. The changes of biochemical indexes indicated that DCZ exposure inhibited lipid synthesis, lipolysis and lipid transport of F0 males and females. In addition, the composition of gut microbes in males and females was significantly changed, which corresponds to changes in biochemical parameters in liver, intestine and serum. Metabolome analysis showed that pathways related to amino acid metabolism, ATP-binding cassette transporters, carbohydrate metabolism, and energy metabolism were downregulated in the gut of males and females. At 120 days post-fertilization, lipid synthesis, lipolysis and lipid transport of F1 males was upregulated; the composition of microbiota and metabolome of the F1males gut was significantly altered. Overall, we found that long-term exposure to low doses of DCZ inhibited the liver-gut axis in males and females, and the liver-gut axis in F1 males were disturbed even in F1 generation without DCZ exposure.

## 1. Introduction

Difenoconazole (DCZ) is a triazole fungicide widely used in agricultural production.^2^ DCZ was frequently detected in the environment and food. In surface waters of rivers in Fujian, China, DCZ was detected at concentrations of 3.9-6.1 ng/L.^3^ The highest detected concentration of DCZ was 0.3 mg/L in surface water of rivers in Kedah, Malaysia.^4^ Furthermore, the highest detected concentration of DCZ was 0.15 μg/L in river surface water in Melbourne, Australia.^5^ In the paddy field water of Changsha, Changchun and Hangzhou, China, the DCZ detection concentration on the day of application was 1.98-2.91 mg/L.^6^ The residual concentration of DCZ in strawberries is 39–96 μg/kg after a 7-day preharvest interval.^1^ Moreover, with the use of DCZ in agricultural products, DCZ may enter the human body through the food chain, thereby causing risks to human health. International Union of Pure and Applied Chemistry showed that the degradation half-lives of DCZ in soil, water sediments and water were 130 days, 1053 days and stable, respectively.^7^ Therefore, more attention should be paid to the risk to animals and humans of DCZ exposure.

In recent years, studies have shown that the host gut microbiota is closely related to the host’s health (energy, immunity, weight, and disease, etc.),^8–10^ and the liver is one of the organs most closely connected and concerned with the gut. Chen et al. (2020) showed that modulation of zebrafish gut microbiota by probiotics could alleviate PFBS-induced hepatic lipid metabolism disorders.^11^ Agricultural fungicide prothioconazole induced liver damage in mice by affecting gut barrier function and disrupting gut microbiota.^12^ A study have shown that DCZ induced disturbance in the gut microbiota and lipid metabolism, and liver damage in zebrafish.^13^ Exposure to DCZ for 30 days resulted in lower levels of cholesterol in the livers of male zebrafish, but no significant changes in females.^4^ After 21 days of DCZ exposure in zebrafish, the concentration of DCZ in females was higher than that in males. Therefore, it is necessary to explore whether DCZ exposure has sex differences in lipid metabolism in the liver-gut axis. In addition, due to the stability of DCZ in the environment, it is necessary to investigate the biological effects of long-term exposure to low-dose DCZ.

To this end, we exposed zebrafish embryos to DCZ (0, 0.1, 1.0 and 10 μg/L) for 120 days and cultured their F1 generation to 120 dpf in the absence of DCZ. After exposure, the growth and development, the composition of gut microbes, the metabolome of gut tissue, and the lipid metabolism between liver-gut-blood of the F0 and F1 generations were analyzed. Eventually, through biochemical, metabolomic, and metagenomics data, we systematically elucidated the effects of long-term exposure to low-dose DCZ in parental zebrafish on liver-gut lipid metabolism in F0 and F1 generations.

## 2. Materials and methods

### 2.1 Chemicals

Difenoconazole (DCZ, CAS NO. 119446-68-3, 96% purity) was purchased from Jiangsu Agrochem Laboratory Co., Ltd (Jiangsu, China) and was dissolved in analytical-grade acetone for the exposure experiments.

### 2.2 Exposure Experiments and Sampling

Zebrafish (wild-type) embryos were obtained and cultivated according to the method published by Mu et al. (2013).^14^ Healthy embryos (about 2 hours postfertilization) were placed in exposure solutions of 0, 0.1, 1.0 and 10 μg/L DCZ for 120 days. The details of exposure process are provided in the Supplementary Information. At 120 dpf, the body length, the weight of body, liver, intestine, gonad and brain of female and male fish were counted, and liver and intestinal samples were collected by dissecting zebrafish on ice. The same tissue from five fish of the same sex as a sample. All samples were flash frozen in liquid nitrogen and stored at −80 °C until analysis.

At 115 dpf, 4 male fish and 4 female fish were transferred separately to 5-L tanks with 2 L exposure solution (the experiment done in triplicate). At 121 dpf, these fish were used for spawning and embryos were collected. The method of spawning experiment was consistent with previous research.^14^ F1 generation embryos were washed six times with reconstituted water, and cultured in a DCZ-free environment until 120 dpf. The breeding process of the F1 generation was the same as that of the F0 generation. At 120 dpf, sampling process of the F1 generation was the same as that of the F0 generation. Since F0 generation in the 10 μg/L DCZ treatment group spawned less and most of the F1 generation in all treatment groups were males, only the F1 generation males in the solvent control, 0.1 and 1.0 μg/L groups were analyzed. Experiments were approved by the Institutional Animal Care and Use Committee of Hebei University.

### 2.3 Biochemical Measurement

Biochemical indicators (total bile acids [TBA], free fatty acids [FFA], triglycerides [TG], low-density lipoprotein cholesterol [LDL-C], high-density lipoprotein cholesterol [HDL-C], and total cholesterol [TC]) in liver, serum, and gut were measured using Nanjing Jiancheng Bioengineering Institute (Nanjing, China) kits, and the total protein concentration was measured using the Bradford protein concentration assay kit from Shanghai Beyotime Biotechnology Co., Ltd (Beijing, China), the detailed detection method was carried out according to the instructions.

### 2.4 16S rRNA gene sequencing

Total genomic DNA samples were extracted using the OMEGA Soil DNA Kit (Omega Bio-Tek, Norcross, GA, USA). The concentration and quality of the extracted DNA were measured using a NanoDrop NC2000 spectrophotometer and agarose gel electrophoresis, respectively. PCR amplification of the V3-V4 region of the bacterial 16S rRNA gene was performed using forward primer 338F (5’-ACTCCTACGGGAGGCAGCA-3’) and reverse primer 806R (5’-GGACTACHVGGGTWTCTAAT-3’). PCR amplicons were purified with Vazyme VAHTSTM DNA Clean Beads (Vazyme, Nanjing, China) and quantified using Quant-iT PicoGreen dsDNA detection kit (Invitrogen, Carlsbad CA, USA). Equal amounts of amplicons were pooled for paired-end 2250 bp sequencing using the Illumina NovaSeq platform and NovaSeq 6000 SP kit (500 cycles) from Shanghai Personal Biotechnology Co., Ltd. (Shanghai, China). Species composition analysis and beta diversity analysis were performed on sequence data using QIIME2 software and R packages (v3.2.0).

### 2.5 Nontargeted Metabolomics Analysis of Intestine

20 mg intestinal tissue of F0 and F1 Generations were respectively taken, and 500 μl of extract solution (methanol: acetonitrile: water = 2: 2: 1 (v/v), containing isotope-labeled internal standard mixture) was added, and the tissue was triturated on ice for 4 min, ultrasound for 5 min. The samples were left at −40°C for 1 h and then centrifuged for 15 min (4°C, 12,000 rpm). Carefully pipette the supernatant via syringe into an injection vial for measurement.

The target compounds were separated using a Waters ACQUITY UPLC BEH Amide (2.1 mm × 100 mm, 1.7 μm) liquid chromatography column under a Vanquish ultra-high performance liquid chromatograph (Thermo Fisher Scientific, USA). Liquid chromatography phase A is an aqueous phase, containing 25 mmol/L ammonium acetate and 25 mmol/L ammonia water, and phase B is acetonitrile. Sample pan temperature: 4 °C, injection volume: 2 μL. The Thermo Q Exactive HFX mass spectrometer is capable of primary and secondary mass spectrometry data acquisition under the control of the control software (Xcalibur, Thermo). After the original data was transformed and annotated by ProteoWizard software, R program package and self built secondary mass spectrometry database, the peaks retained after preprocessing were subjected to univariate statistical analysis (Student’s t test) and orthogonal partial least squares discrimination analysis (OPLS-DA); the differential metabolites were screened with the P value of Student’s t test less than 0.05 and the criteria of variable importance in the projection (VIP) value of the OPLS-DA model greater than 1.0; then the differential metabolites were subjected to hierarchical clustering analysis and KEGG metabolic pathway analysis.

### 2.6 Gene Expression Analysis

The liver of F0 and F1 Generations were sampled to analyze the expression levels of genes related to the lipid metabolism. All primers of target genes were adopted from previous studies,^11, 15^ and listed in Table S1. Detailed steps for gene expression analysis are provided in Supplementary Information.

### 2.7 Statistical Analyses

A one-way analysis of variance (ANOVA), followed by a Fisher’s LSD post-hoc test, was performed in SPSS (version 16.0, IBM, USA). *P*\*<0.05 were considered statistically significant.

## 3. Results

### 3.1 Effects of DCZ on Growth and Development of F0 and F1 Generations

As shown in Table 1, compared with the control group, the body weight of F0 female fish significantly increased in the 0.1 μg/L DCZ exposure group, and significantly decreased in the 10 μg/L DCZ exposure group. In all treatment groups, the body length of F0 female fish significantly increased, and CF decreased. For the tissues, BSI and GSI of F0 female fish were significantly decreased in the 1.0 μg/L DCZ exposure group. Overall, the growth and development of F0 females was inhibited after DCZ exposure. Likewise, the growth and development of F0 male fish was inhibited after DCZ exposure (Table 1). Compared with the control group, the body weight and CF of F0 male fish were significantly reduced at 1.0 and 10 μg/L DCZ. For the tissues of male fish, HSI was significantly increased at 0.1 μg/L DCZ; GSI was significantly decreased at 0.1 and 10 μg/L DCZ, while in the 1.0 μg/L DCZ treatment group, GSI was significantly increased, which may be caused by the significant decrease in the body weight of F0 male fish in the 1.0 μg/L DCZ treatment group.

**Table 1.** Changes in the growth and development parameters of F0 generation fish.

| Parameters | Control | 0.1 µg/L | 1.0 µg/L | 10 µg/L |
| --- | --- | --- | --- | --- |
| Female |  |  |  |  |
| Body Weight (mg) | 418.3 ±6.6 | 458.3 ±3.7* | 423.1 ±21.0 | 380.9 ±16.7* |
| Body length (mm) | 36.1 ±0.6 | 37.5 ±0.2* | 38.2 ±0.7* | 38.1 ±0.4* |
| CF | 0.89 ±0.06 | 0.87 ±0.01 | 0.76 ±0.01* | 0.69 ±0.03* |
| BSI | 1.14 ±0.03 | 1.18 ±0.15 | 1.00 ±0.20* | 1.43 ±0.32 |
| HSI | 2.63 ±0.48 | 2.75 ±0.15 | 2.86 ±0.16 | 2.95 ±0.50 |
| ISI | 3.41 ±0.23 | 2.99 ±0.16 | 3.17 ±0.23 | 3.79 ±0.62 |
| GSI | 8.90 ±1.4 | 12.5 ±1.4* | 11.1 ±1.5 | 5.88 ±1.73* |
| Male |  |  |  |  |
| Body Weight (mg) | 335.6 ±7.2 | 326.2 ±9.6 | 299.2 ±10.38* | 292.4 ±27.9* |
| Body length (mm) | 35.7 ±0.3 | 35.9 ±0.1 | 35.2 ±0.5 | 36.1 ±0.6 |
| CF | 0.74 ±0.03 | 0.70 ±0.01 | 0.68 ±0.00* | 0.62 ±0.04* |
| BSI | 1.70 ±0.07 | 1.83 ±0.27 | 1.69 ±0.16 | 1.62 ±0.54 |
| HSI | 0.70 ±0.09 | 1.25 ±0.06* | 1.13 ±0.26 | 1.10 ±0.40 |
| ISI | 2.38 ±0.02 | 2.24 ±0.18 | 2.39 ±0.42 | 2.56 ±0.27 |
| GSI | 1.17 ±0.09 | 0.87 ±0.07* | 1.45 ±0.15* | 0.64 ±0.22* |

Note: Condition factor (CF)=body weight (mg)×100/body length (mm)<sup>3</sup>; brain-somatic index (BSI)=brain weight (mg)×100/body weight (mg); hepatosomatic index (HSI)=hepatic Weight (mg)×100/body weight (mg); Intestinal index (ISI)=intestinal weight (mg)×100/body weight (mg); gonadosomatic index (GSI)=gonad weight (mg)×100/body weight (mg). Values are shown as the mean ± SD of three replicates per treatment (\**p* < 0.05).

As shown in Table S2, parental exposure to DCZ did not significantly alter the growth and development of F1 male zebrafish at 120 dpf.

### 3.2 Effects of DCZ on Lipid metabolism of F0 and F1 Generations

As shown in Table S3, compared with the control group, the liver TC content of F0 female fish was significantly reduced in all treatment groups; 10 μg/L DCZ significantly reduced the contents of liver TG, liver FFA and serum TC of F0 female fish; 1.0 and 10 μg/L DCZ significantly reduced TBA content in liver, serum and intestine of F0 female fish, and LDL-C content, HDL-C content and TG content in F0 female fish serum. From Table S4, it can be seen that the expression level of genes (such as *lipca*, *cyp8b1*, *cyp27a1*, *hsd3b7*, *cs*, *cpt2*, *pparg* and *apoba*, etc.) in the liver of F0 female fish were significantly down-regulated after DCZ exposure, the expression level of a few genes (such as *lipea*, *cyp7a1* and *cpt1* at 1.0 μg/L DCZ) was up-regulated.

For F0 male fish, most of the biochemical indicators related to lipid metabolism were significantly inhibited (Table S5). Compared with the control group, the contents of TC and TBA in the liver and the contents of HDL-C, TC and TBA in the serum were significantly decreased in all DCZ treatment groups; 1.0 and 10 μg/L DCZ significantly reduced FFA content in the liver, LDL-C content in the serum and TBA content in the intestine; 1.0 and 10 μg/L DCZ significantly reduced TG content in the serum and TG content in the liver, respectively. In addition, Table S6 showed that the expression level of genes (such as *cyp7a1*, *cyp8b1*, *cpt2*, *mcad*, *acox1*, *pparg*, *mttp* and *apoba,* etc.) was down-regulated in the liver of F0 male fish after DCZ exposure, the expression level of a few genes (such as *cs*, *aclya* and *srebp1* in the 10 μg/L DCZ group) was up-regulated.

For F1 male fish (Table S7), compared with the control group, TBA content in the liver, serum and intestine significantly increased at 1.0 μg/L DCZ, and TG content in the serum was significantly decreased at 0.1 and 1.0 μg/L DCZ. In addition, Table S8 showed that the expression level of genes (such as, *lipca*, *cyp7a1*, *cyp8b1*, *cpt2*, *mcad*, *dgat2*, *acox2*, *srebp1*, *pparg*, *mttp*, and *apoba,* etc.) was up-regulated in the liver of F1 male fish.

### 3.3 Effects of DCZ on Gut Microbiota of F0 and F1 Generations

The principal coordinate analysis (PCoA) showed that DCZ-treated group and control group can be clearly separated (Figure S1), indicating that DCZ exposure significantly altered the gut microbiota composition of F0 female fish, F0 male fish and F1 male fish.

At the phylum and genus level, compared with the control group, there were significant differences in the gut microbiota composition of F0 female and male fish in the DCZ-treated group (Figure 2 and 3). The abundance of *Fusobacteria* in F0 female fish was significantly increased at 0.1 μg/L DCZ, while 1.0 and 10 μg/L DCZ significantly reduced the abundance of *Fusobacteria* (Figure 2B). The abundance of *Bacteroidetes* and *Firmicutes* of F0 male fish was significantly increased at 0.1 or 1.0 μg/L DCZ (Figure 2D). The abundance of *Rhodobacter* of F0 female fish was significantly decreased in 0.1 μg/L DCZ-treated group, but significantly increased at 10 μg/L DCZ (Figure 3B). The abundance of *Cetobacterium* of F0 female fish was significantly increased at 0.1 μg/L DCZ, but significantly decreased at 1.0 and 10 μg/L DCZ (Figure 3B). Figure 3D showed that the abundance of *Rhodobacter* of F0 male fish was significantly decreased at 0.1 or 1.0 μg/L DCZ, but increased at 10 μg/L DCZ.

**Figure 1.**
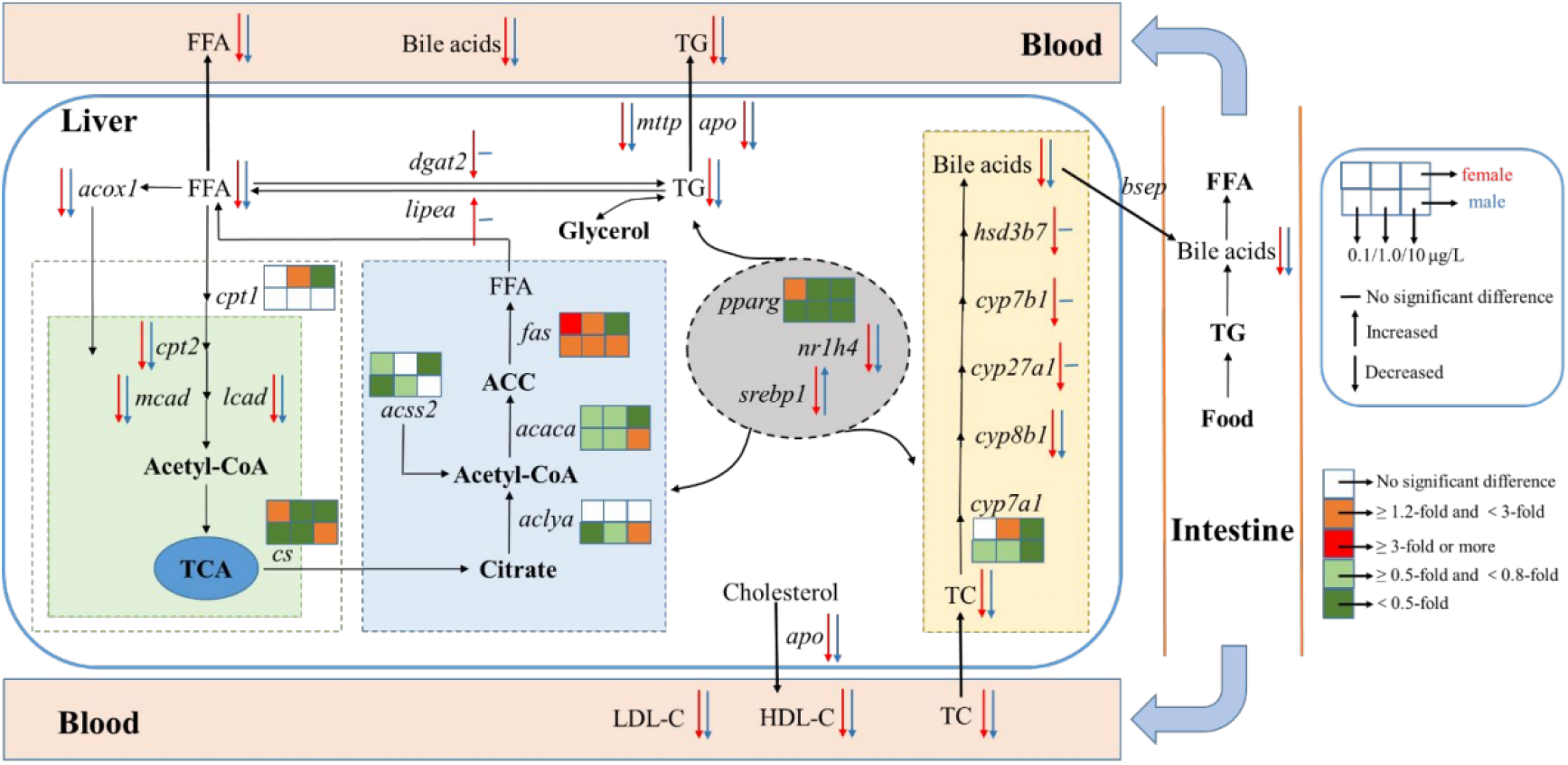
Integrative map of lipid metabolism activity in intestine, blood, and liver tissues from F0 female and male zebrafish after a 120-day exposure to DCZ (0.1, 1.0, and 10 μg/L). Values are presented as the mean of three replicates (n = 3). Italic characters indicate gene transcriptions. Non-bold characters indicate the metrics tested in this study.

**Figure 2.**
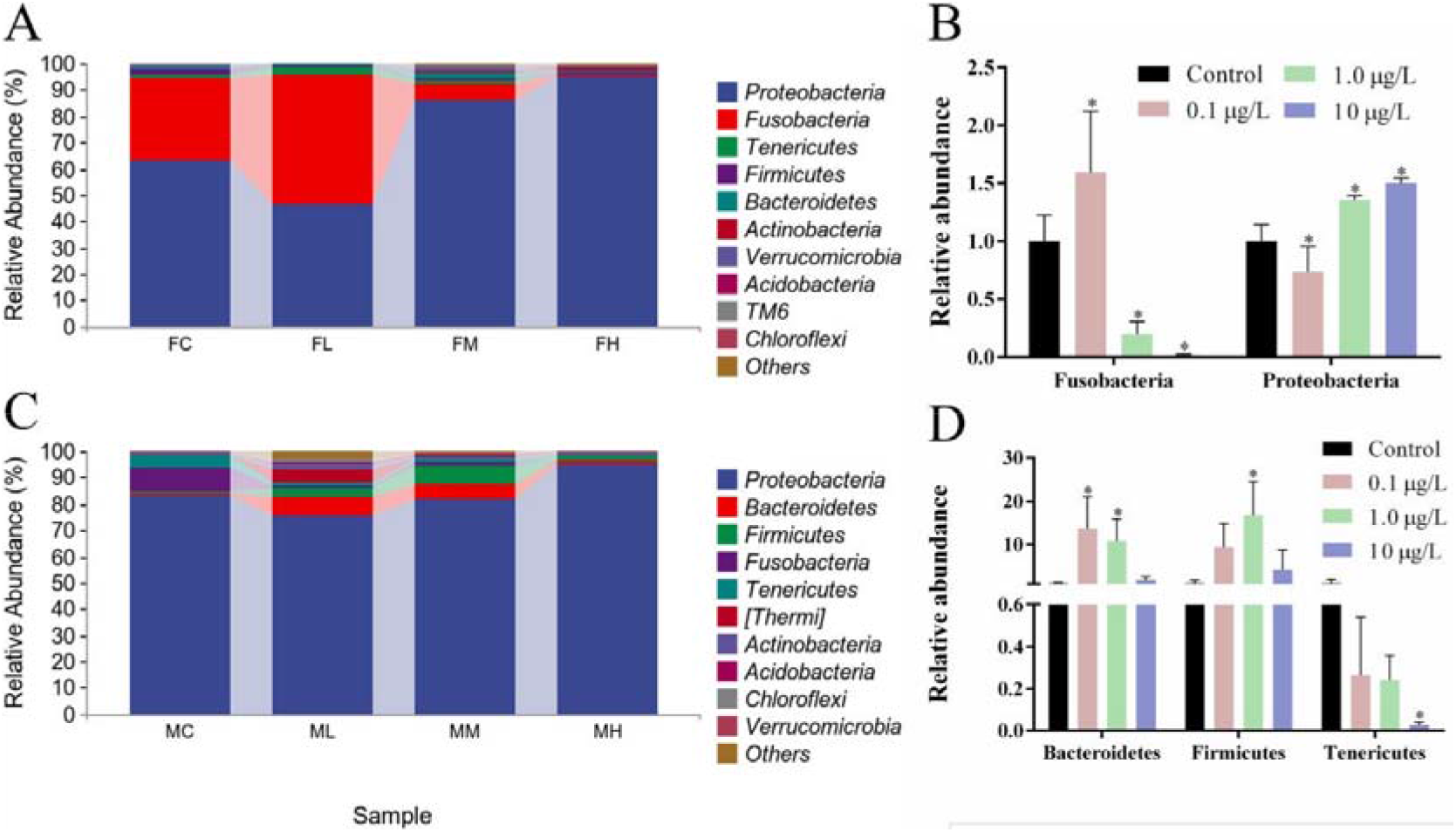
The effect of DCZ exposure on the phylum level of gut microbiota in F0 generation zebrafish. A: The composition of the gut microbiota of F0 generation female fish at the phylum level; B: The main representative microbiota changes at the phylum level of F0 female fish; C: The composition of gut microbiota of F0 generation male fish at the phylum level; D: The main representative microbiota changes at the phylum level of F0 male fish. FC: F0 female fish-control; FL: F0 female fish-0.1 μg/L DCZ; FM: F0 female fish-1.0 μg/L DCZ; FH: F0 female fish-10 μg/L DCZ; MC: F0 male fish-control; ML: F0 male fish-0.1 jrg/L DCZ; MM: F0 male fish-1.0 μg/L DCZ; MH: F0 male fish-10 μg/L DCZ. (\**p*< 0.05).

**Figure 3.**
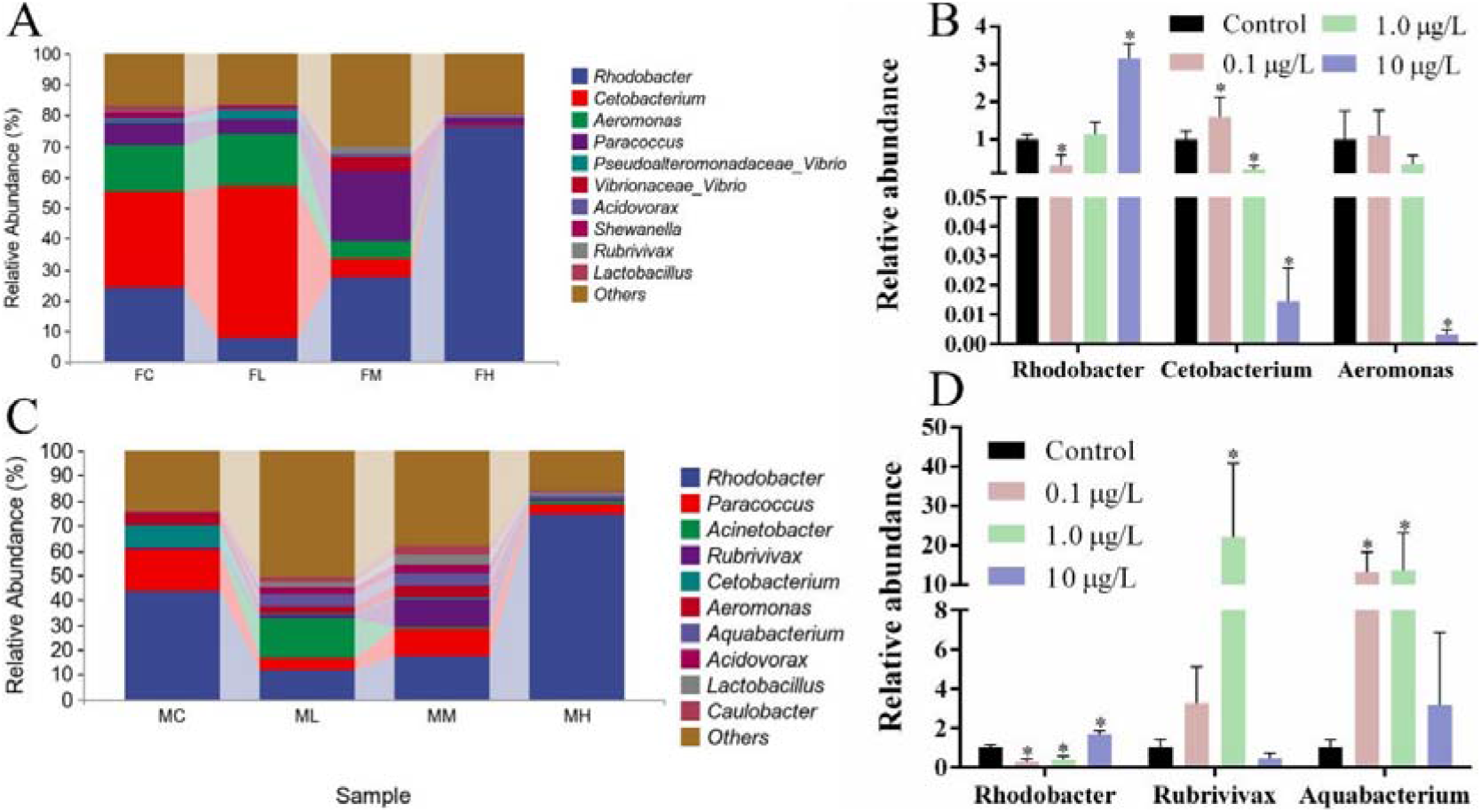
The effect of DCZ exposure on the genus level of gut microbiota in F0 generation zebrafish. A: The composition of the gut microbiota of F0 generation female fish at the genus level; B: The main representative microbiota changes at the genus level of F0 female fish; C: The composition of gut microbiota of F0 generation male fish at the genus level; D: The main representative microbiota changes at the genus level of F0 male fish. (\**p*< 0.05).

For F1 male fish (Figure 4), compared with the control group, the gut microbiota composition in the DCZ-treated group was significantly changed at both phylum and genus levels. At the phylum level (Figure 4C), the abundance of *Proteobacteria* was significantly decreased at 1.0 μg/L DCZ; the abundance of *Actinobacteria* and *Fusobacteria* was significantly increased at 0.1 or 1.0 μg/L DCZ. At the genus level (Figure 4D), the abundance of *Rhodobacter* was significantly increased at 0.1 or 1.0 μg/L DCZ; the abundance of *Acinetobacter* was significantly decreased at 0.1 and 1.0 μg/L DCZ.

**Figure 4.**
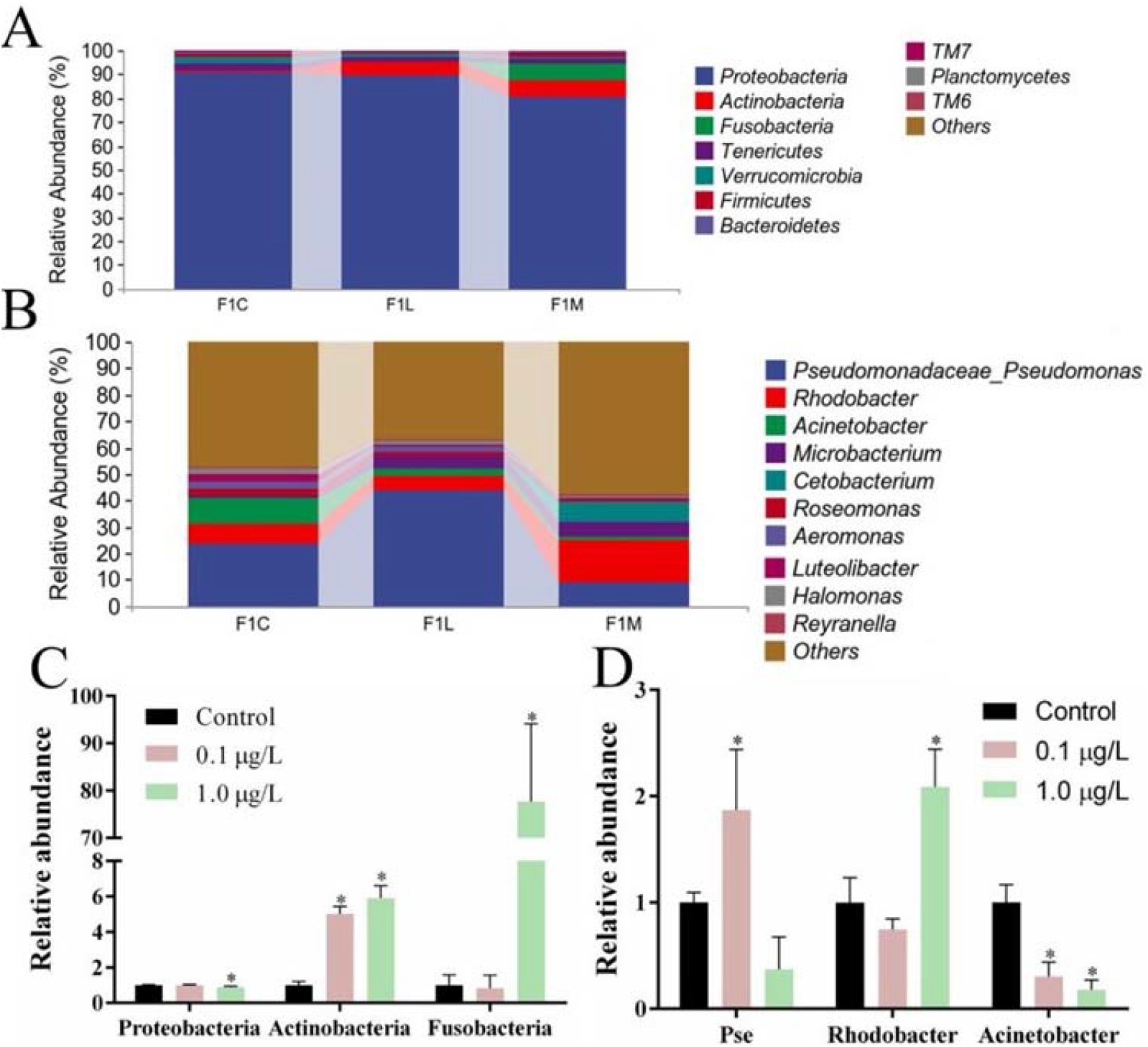
The effect of DCZ exposure on the phylum and genus level of gut microbiota in F1 generation zebrafish. A: The composition of the gut microbiota of F1 generation male fish at the phylum level; B: The composition of gut microbiota of F1 generation male fish at the genus level; C: The main representative microbiota changes at the phylum level of F1 male fish; D: The main representative microbiota changes at the genus level of F1 male fish. F1C: F1 male fish-control; F1L: F1 male fish-0.1 μg/L DCZ; F1M: F1 male fish-1.0 μg/L DCZ. (\**p*< 0.05).

### 3.4 Effects of DCZ on Intestine Metabolomes of F0 and F1 Generations

As shown in Figure S2A\C\E, the DCZ-treated group and the control group could be clearly separated, indicating that DCZ exposure significantly altered the intestinal metabolism of F0 female fish, F0 male fish and F1 male fish. For the gut of F0 female fish (Figure S2B), there were 803 down-regulated differential metabolites and 524 up-regulated differential metabolites at 10 μg/L DCZ; for the gut of F0 generation male fish (Figure S2D), there were 2086 down-regulated differential metabolites and 572 up-regulated differential metabolites at 10 μg/L DCZ; for the gut of F1 male fish (Figure S2F), there were 4109 down-regulated differential metabolites and 812 up-regulated differential metabolites at 1.0 μg/L DCZ. The heat map also further indicated that the gut metabolism of F0 females, F0 males and F1 males was disturbed after DCZ exposure (Figure S3-S5). KEGG pathway analysis of differential metabolites showed that DCZ exposure mainly affected the pathways related to amino acid metabolism, ATP-binding cassette transporter (ABC transporter), and glycolipid metabolism in the intestine of F0 female and male fish and F1 male fish (Figure 5).

**Figure 5.**
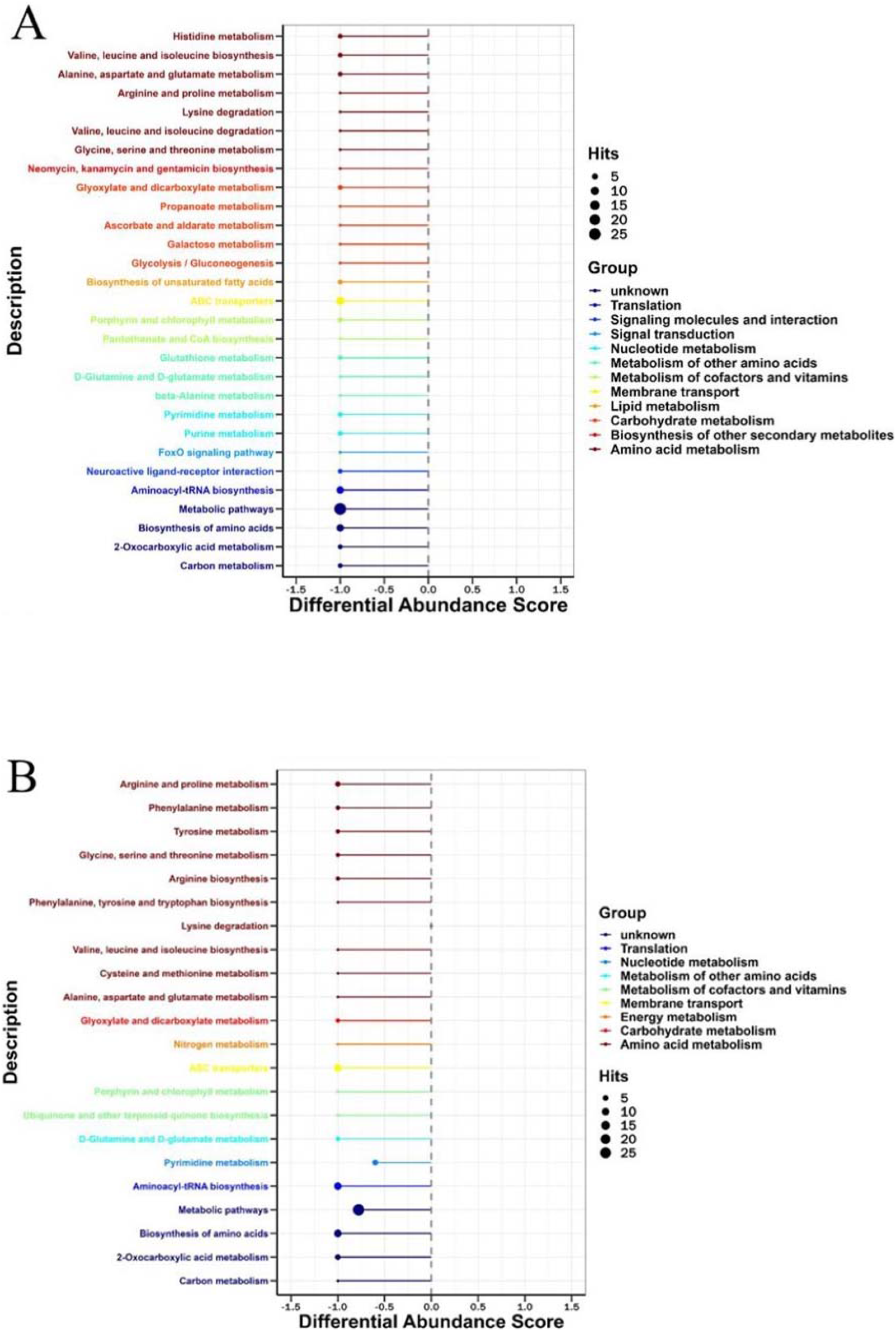
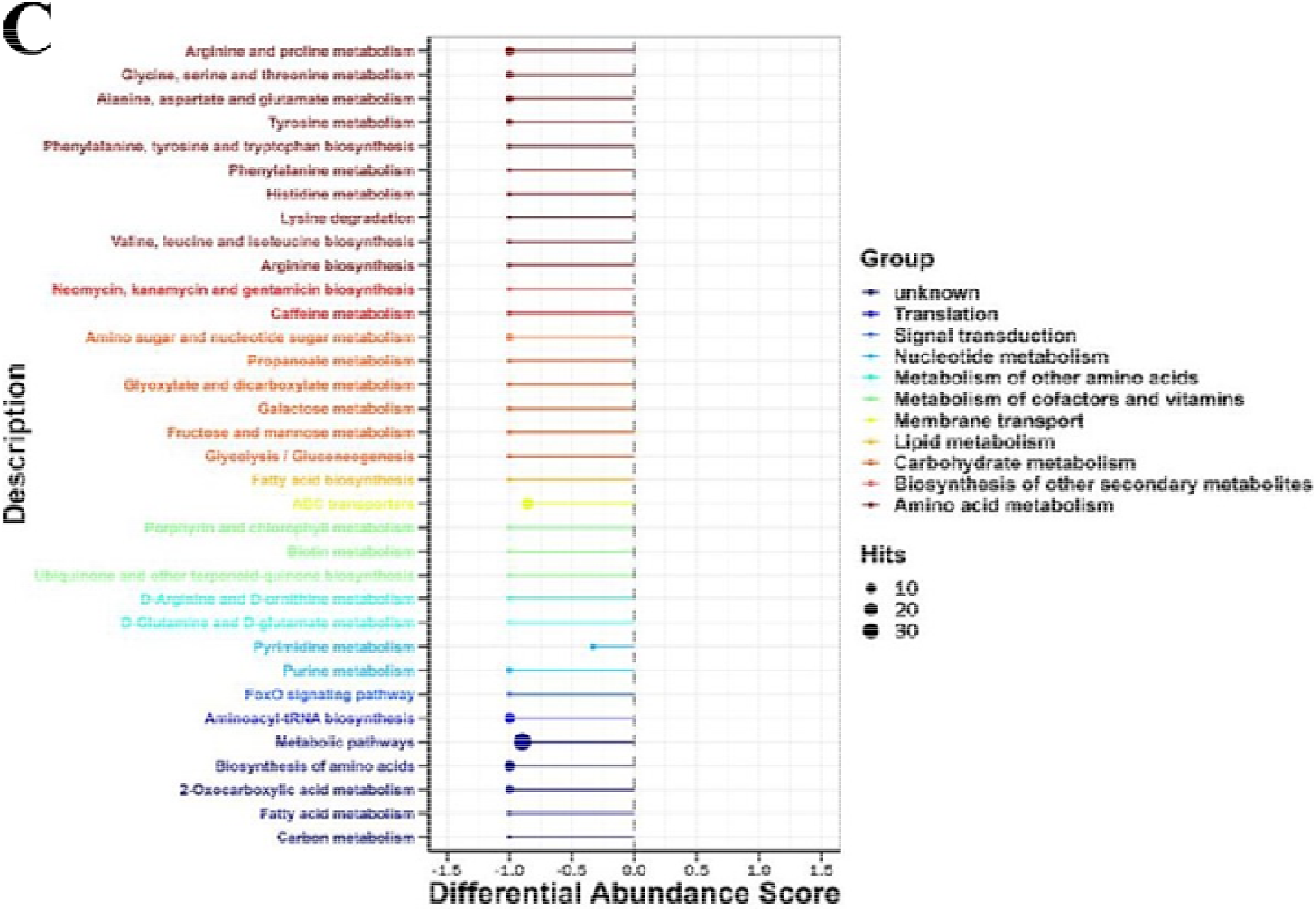
KEGG pathway analysis of differential metabolites in the gut of F0 and F1 zebrafish. A: KEGG pathway analysis of differential metabolites in the gut of F0 female zebrafish; B: KEGG pathway analysis of differential metabolites in the gut of F0 male zebrafish; C: KEGG pathway analysis of differential metabolites in the gut of F1 male zebrafish. If the value of the line segment is normal, it means that the entire pathway was up-regulated; otherwise, it means that the pathway was down-regulated.

## 4. Discussion

In this study, it was found that the growth and development of F0 generation female and male fish at 120 dpf were significantly affected after full-cycle exposure to DCZ, including decreased body weight of male and female fish, increased body length of female fish, reduced CF, and GSI at 1.0 and 10 μg/L DCZ, while female fish at 0.1 μg/L DCZ had significantly higher body weight and GSI, and HSI of male fish at 0.1 μg/L DCZ was significantly increased. Therefore, low concentrations of DCZ may induce self-defense and stress responses in fish, thereby promoting growth and development. Teng et al. (2017) found that exposure to DCZ (5, 50, and 500 μg/L) for 7 days significantly altered the expression level of genes related to the growth hormone endocrine (GH) axis in adult zebrafish, and there are sex differences in the effect of DCZ on the axis.^16^ In this study, the body length of F0 female fish was significantly increased while the body length of male fish was not significantly change, which may be caused by the sex differences in the effect of DCZ on the GH axis. And the effect of DCZ on the GH axis may be one of the reasons for the abnormal growth and development of F0 female and male fish. In addition, our results showed that the effect of DCZ on liver-gut lipid metabolism was also one of the reasons for abnormal growth and development of F0 male and female fish.

Lipid metabolism plays a very important role in energy supply and growth and development.^17–19^ This study comprehensively explored fundamental metabolic pathways associated with lipid metabolism in the liver, blood, and gut, including gut digestion, bile acid synthesis and excretion, blood transport, nuclear signaling, fatty acid biosynthesis and β-oxidation, and TG biosynthesis. In the current study, the lipid metabolism processes that were mainly affected in F0 generation male and female fish were fatty acid biosynthesis and β-oxidation, TG biosynthesis, nuclear signaling, and bile acid synthesis (Figure 1). Fatty acids are the most energy-producing substances in animals.^20^ Fatty acid biosynthesis and β-oxidation directly affect energy supply.^11^ The function of *aclya* (ATP-citrate lyase a) is to cleave citrate to form acetyl-CoA, thereby providing a substrate for fatty acid synthesis.^21^ In this study, the expression level of *aclya* in the liver of F0 generation male and female fish was significantly down-regulated, which led to the reduction of FFA content in the liver. Moreover, the reduction of FFA content in the liver further inhibited fatty acid β-oxidation, and the expression levels of *cpt2*, *cpt1*, *mcad*, *lacd* and *acox1* related to the fatty acid β-oxidation process were significantly down-regulated in the liver of F0 generation male and female fish (Figure 1). Diacylglycerol O-acyltransferase 2 (dgat2) catalyzes the conversion of fatty acids and glycerol to triglycerides, which is the final step in TG biosynthesis.^22^ The expression level of *dgat2* was also significantly down-regulated due to the reduction of FFA content in the liver of F0 generation male and female fish. FFA can bind to and activate the nuclear receptor peroxisome proliferator-activated receptor (PPAR), which plays a key role in regulating lipid metabolism.^23, 24^ The expression level of *ppar* in the liver of F0 generation female and male fish was significantly down-regulated, which may be induced by the reduction of FFA content. Similarly, the expression level of the nuclear receptor *farnesoid X receptor* (*nr1h4*), which is associated with bile acid homeostasis,^25^ was significantly down-regulated. In addition, the expression level of *apolipoprotein* (*apoa1a*, *apob1* and *apoc1*), which are involved in the mutual transport of cholesterol between the liver and peripheral tissues,^26–28^ were also significantly down-regulated. Consistently, the levels of LDL-C and HDL-C in the blood of F0 generation male and female fish were also significantly reduced. The metabolic pathway of ABC transporter in the gut of F0 generation male and female fish was also down-regulated (Figure 5). Binding of ABC transporters to LDL-C and HDL-C mediates cholesterol transport across cell membranes.^26^ Bile acids are synthesized in the liver using cholesterol as a substrate and act as emulsifiers in the gut to facilitate lipid absorption.^29^ In this study, cholesterol, the raw material for bile acid synthesis, was significantly reduced in the liver of F0 male and female fish, and the expression levels of genes involved in the bile acid synthesis pathway (for example, *cyp8b1*, *cyp27a1*, *cyp7b1,* and *hsd3b7)* significantly down-regulated (Figure 1). Morrison et al. (2014) found that the imidazole fungicide ketoconazole and the triazole fungicide propiconazole inhibited sterol 14α-demethylase (CYP51) in zebrafish through homology simulation and molecular docking techniques,^30^ and CYP51 is a key enzyme in the cholesterol synthesis pathway.^31^ Therefore, inhibition of CYP51 by DCZ inhibited the bile acid synthesis pathway in the liver of F0 male and female fish. Consistently, DCZ exposure significantly reduced the concentration of TBA in the gut of F0 generation male and female fish, which is bound to affect the absorption and metabolism of lipids in the gut. It was mentioned that both FFA and TG contents in the liver of F0 generation male and female fish were significantly reduced in the DCZ exposure group.

The gut microbiota is known to tightly regulate the physiological functions of host organisms, especially nutrient digestion and metabolism.^32–35^ In this study, the gut microbiota of F0 generation male and female fish was affected. For the gut of F0 female fish, the abundance of *Cetobacterium* was significantly down-regulated at 1.0 and 10 μg/L DCZ. *Cetobacterium* is important to the fish gut microbiome and is involved in promoting vitamin B12 synthesis.^36, 37^ Vitamin B12 plays an important role in lipid metabolism.^38, 39^ In the 0.1 μg/L DCZ group, the abundance of *Cetobacterium* in F0 male fish was significantly up-regulated, which was a compensatory response of the fish itself and corresponded to the increased body weight and gonadal index of F0 generation zebrafish in the 0.1 μg/L DCZ group. Previous study have shown that decreased abundance of *Bacteroides* and increased abundance of *Firmicutes* increased body weight in mice.^40^ The increased abundance of *Bacteroides* and *Firmicutes* in the gut of F0 males may have induced changes in their body weight. *Rhodobacter* boosts bile acid production in the gut.^41^ In the 0.1 or 1.0 μg/L DCZ group, the abundance of *Rhodobacter* in F0 male and female fish was significantly reduced, which further exacerbated the reduction in intestinal bile acid content, while in the 10 μg/L DCZ group, the significantly increased abundance of *Rhodobacter* in F0 generation male and female fish may be a compensatory response of the body. Pathways related to the metabolism of cofactors and vitamins, lipid metabolism and carbohydrate metabolism in F0 male and female fish were down-regulated (Figure 5A/B), which further reflect the effects of DCZ on gut microbiota and hepatic lipid metabolism.

The results of this study showed that after full-cycle exposure of F0 zebrafish to DCZ, even if the F1 generation was not exposed to DCZ, the liver-gut axis of the F1 generation was still affected, but this was not sufficient to affect growth and development of F1 generation at 120 dpf. Contrary to F0 generation of male and female fish, the expression level of genes related to lipid metabolism (such as, intestinal digestion, bile acid synthesis and excretion, blood transport, nuclear signaling, fatty acid biosynthesis and β-oxidation, and TG biosynthesis, etc.) in the liver of F1 male fish were significantly up-regulated. We speculate that the up-regulation of lipid metabolism pathway may be a compensatory response to the congenital deficiency of F1 males. A previous study showed that the growth and development of F1 generation embryos/larvae were significantly affected in adult zebrafish exposed to DCZ for 21 days.^42^ And we also observed abnormal growth and development, increased apoptosis and decreased activity of enzymes related to energy metabolism in the embryo/larvae stage of F1 generation. At 120 dpf, the reduction of TG content in serum of F1 male fish (Table S7) and the down-regulation of pathways related to cofactor and vitamin metabolism, lipid metabolism, ABC transporters, and carbohydrate metabolism in the gut of F1 male fish (Figure 5C) was a manifestation of congenital deficiency. Under the same culture conditions as the F1 generation of the control group, the F1 generation of the exposed group must compensate for the congenital deficiency through its own compensation response. In the liver, serum and intestine of F1 male fish, the content of TBA was significantly increased to promote the digestion and absorption of lipids. *Rhodobacter* boosts bile acid production in the gut.^41^ The abundance of *Rhodobacter* was significantly increased in F1 males exposed to DCZ, which may be one of the reasons for the increase in intestinal bile acid content.

In summary, the present study shows that long-term exposure to DCZ at ambient concentrations inhibited pathways related to lipid metabolism, including fatty acid biosynthesis and β-oxidation, TG biosynthesis, nuclear signaling, and bile acid synthesis, in the liver, blood, and gut of F0 male and female fish. The composition of gut microbiota *(Cetobacterium, Rhodobacter* and *Firmicutes,* etc.) in F0 generation male and female fish was significantly changed, and the pathways related to cofactor and vitamin metabolism, glycolipid metabolism and ABC transporter were downregulated in the gut. In addition, DCZ exposure inhibited the growth and development of F0 generation male and female fish, such as decreased body weight, decreased growth factors and decreased gonadal index. In conclusion, the disruption of F0 liver-gut axis by DCZ inhibited the growth and development of parental male and female fish. Full-cycle exposure of the parent to DCZ without further exposure in the F1 generation still had an effect on the liver-gut axis of the F1 generation male fish, which was not sufficient to alter the growth and development of the F1 generation male fish at 120 dpf.

## Acknowledgments

This work was supported by the Advanced Talents Incubation Program of Hebei University (No. 050001-521000081469).

## Conflict of interest

The authors declare that they have no known competing financial interests or personal relationships that could have appeared to influence the work reported in this paper.

## Graphic for Table of Contents

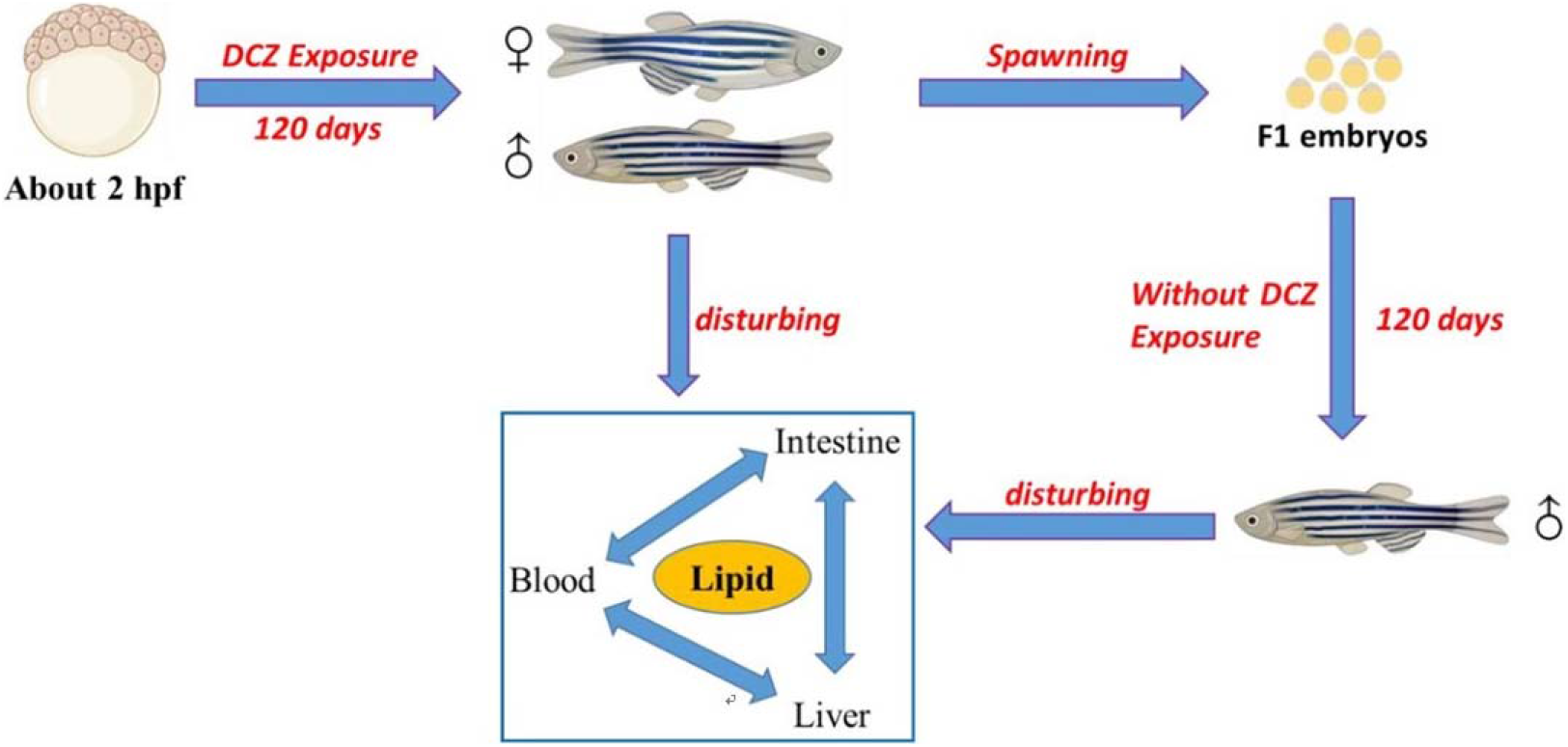

## Notes

### Competing Interest Statement

The authors have declared no competing interest.

