## Supplementary Information for "Effects of Full-cycle Exposure to Difenoconazole in Parental Zebrafish on the Liver-gut Axis of F0 and F1 Generations"

**Summary of Supporting Information**

**Methods:** Exposure Experiments, Feeding procedure, Gene Expression Analysis.

**Tables and Figures:**

Table S1. Primer pair sequences for gene transcriptional analysis.

Table S2. Changes in the growth and development parameters of F1 male zebrafish.

Table S3. Biochemical alterations in liver, blood and intestine tissues of F0 female zebrafish.

Table S4. Gene transcriptional changes involved in hepatic lipid metabolisms of F0 female zebrafish.

Table S5. Biochemical alterations in liver, blood and intestine tissues of F0 male zebrafish.

Table S6. Gene transcriptional changes involved in hepatic lipid metabolisms of F0 male zebrafish.

Table S7. Biochemical alterations in liver, blood and intestine tissues of F1 male zebrafish.

Table S8. Gene transcriptional changes involved in hepatic lipid metabolisms of F1 male zebrafish.

Figure S1. Principal coordinate analysis (PCoA) of the gut microbiome of F0 and F1 generations.

Figure S2. OPLS-DA plot and volcano plot of DCZ exposure on intestinal metabolism of F0 and F1 generations.

Figure S3. Heat map of differential metabolites in F0 female fish gut

Figure S4. Heat map of differential metabolites in F0 male fish gut.

**References**

#### METHODS

**Exposure Experiments**. Zebrafish (wild-type AB-strain) embryos were obtained and cultivated according to the method published by Mu et al. (2013). Healthy embryos (about 2 hours post-fertilization (hpf)) were placed in exposure solutions of 0, 0.1, 1.0 and 10 μg/L DCZ for 120 days. Exposure concentrations were selected based on the detected concentration (0.15 μg/L) of DCZ in surface water in Australia (Schafer et al., 2011). Both blank control and solvent control were set. Except for the blank control, the acetone content in the other treatment groups was 0.0005%. 70 embryos were put into a 1-L beaker containing 0.5 L exposure solution, and the experiment done in triplicate (Three beakers per treatment group, each beaker containing 70 embryos). At 12 days post-fertilization (dpf), larvae in the beakers were transferred to 5-L tanks with 2 L exposure solution per tank (Three tanks per treatment group, each tank containing 68-70 larvae). At 21 dpf, fish in the tanks were moved to 20-L tank with 5 L exposure solution per tank (Three tanks per treatment group, each tank containing 60-64 zebrafish). At 60 dpf, the exposure solution in each 20-L tank changed from 5 L to 15 L until 120 dpf (Three tanks per treatment group, each tank containing 60-64 zebrafish). From about 2 hpf to 21 dpf, the mortality rate of the fish in each tank was 9-14%, and during the period from 21 dpf to the end of the experiment, no dead individuals were observed. The details of feeding process are provided in the Supplementary Information.

In the beginning four days, exposure solutions were renewed every 24 h. During the period from 5 dpf to 21 dpf, exposure solutions were renewed half an hour after feeding, and during the period from 21 dpf to the end of the experiment, exposure solutions were renewed every two days.

**Feeding procedure.** Zebrafish larvae were fed daily as follows: 5-11 dpf, zebrafish larvae were fed with commercial larvae feed (GEMMA Micro 75, Skretting, USA) four times per day (0.05 g/tank); 12-20 dpf, zebrafish larvae were fed with commercial larvae feed (0.05 g/tank) and brine shrimp (0.1 g/tank) three times per day; 21-120 dpf, zebrafish larvae were fed with brine shrimp three times per day, the feeding amount increased with growth (the feeding volume was determined as the amount which could be eaten in 10-15 min).

**Gene Expression Analysis.** The liver of F0 and F1 Generations were sampled to analyze the expression levels of genes related to the lipid metabolism. Total RNA was extracted from zebrafish liver, following the TRIzol reagent protocol (Tiangen Biotech, Beijing, China). The concentration of RNA was measured by a Nano Drop 2000c spectrophotometer (Thermo Scientific, Wilmington, DE). The banding pattern was on a 1% agarose gel electrophoresis. The purity was assessed by the A260/A280 ratio. The cDNA was then synthesised via a quant RTase kit (Tiangen Biotech, Beijing, China) in accordance with the manufacturer's recommendations. Real-time quantitative polymerase chain reaction (RT-qPCR) were performed with the ABI 7500 q-PCR system (Applied Biosystems). SYBR Green PCR Master Mix reagent kits (Tiangen

Biotech) were used for the quantification of gene expression. A 20μL reaction system was conducted according to the manufacturer's instructions. The PCR amplification procedure was as follows: denaturation for 15 min at 95 °C, followed by 40 cycles of amplification of 10s at 90°C, 20s at 60 °C, and 32s at 72°C. All primers of target genes were based on previous studies (Qian et al., 2021; Chen et al., 2020), and listed in Table S1. The housekeeping gene *β*-actin was used as an internal control. The relative transcription levels of the target genes were calculated using the $2^{-\Delta\Delta Ct}$ method (Livak and Schmittgen, 2001).

### TABLES AND FIGURES

**Table S1.** Primer pair sequences for gene transcriptional analysis.

| Target gene | Full name | Forward primer (5'-3') | Reverse primer (3'-5') | References |
|---|---|---|---|---|
| β-actin | Beta-actin | TGGACTCTGGTGATGGTGTGAC | GAGGAAGAAGAGGCAGCGGTTC | (Qian et al., 2021) |
| lipca | lipase hepatic a | GGCTTATCTTTTGGGGCTTC | CAGTGTGTGAGGCTGGAAGA | (Chen et al., 2020) |
| lpl | lipoprotein lipase | CTGGCCTTCTCACCAAACAT | GCCTTTGAATCCCAATGCTA | (Chen et al., 2020) |
| lipea | hormone sensitive lipase a | CCCTCTGCTGCCTTTAAGTG | GTTGCAGAGGCTGTTGATGA | (Chen et al., 2020) |
| cyp7a1 | cholesterol 7-alpha-monooxygenase | GATCTTCCCAGCTCTGATCG | GCAGAGTGTTGGCTTGTGAA | (Chen et al., 2020) |
| cyp8b1 | sterol 12α-hydroxylase | GTGGTCAAAGAGGCAAGAGC | ACACTTTGCCATTCCCAGAC | (Chen et al., 2020) |
| nr1h4 | nuclear receptor subfamily 1, group H, member 4 (farnesoid X receptor) | CACAACAAACATCGCATTCC | GCTGAAGACTTGGGCTGAAC | (Chen et al., 2020) |
| cyp27a1 | sterol 27-hydroxylase | TCCAAAGGACTCGCTTCAGT | GCGTCTCGAAGAGAATGGAC | (Chen et al., 2020) |
| cyp7b1 | oxysterol 7α-hydroxylase | CCGAACAGCTCTTTTTCAGG | TTTTTCTCTCCGAGCACGTT | (Chen et al., 2020) |
| hsd3b7 | 3β-hydroxy-Δ5-C27-steroid oxidoreductase | CTCTGCAGGAACATCCCAAT | TGATCCACAGCATCCACACT | (Chen et al., 2020) |
| bsep | bile salt export pump | AACGCTGAAGATGCTGACCT | GTCATCATGCCGTACACCAG | (Chen et al., 2020) |
| cs | citrate synthase | AGCGCTGCTATGAATGGTCT | GTGCTTGAGGGCAAACTCTC | (Chen et al., 2020) |
| aclya | ATP-citrate lyase a | CCTGTGCCACTCTCTTCTCC | CACACCCTTGGCTACCAGAT | (Chen et al., 2020) |
| acaca | acetyl-CoA carboxylase alpha | GCAAGTGTGGTTCCCTGATT | ATACACCAGAACCGGCTGTC | (Chen et al., 2020) |
| fas | fatty acid synthase | GCACCGGTACTAAGGTTGGA | ACACAACCGACCATCTGTCA | (Chen et al., 2020) |
| cpt2 | carnitine palmitoyltransferase 2 | GTGCCTGATGGTTTTGGAGT | CAATGAATGGGGTTTCCATC | (Chen et al., 2020) |
| cpt1 | carnitine palmitoyltransferase 1 | GTGCGGTCTTGCACTACAGA | TGGACAGTCTCCAAGGCTCT | (Chen et al., 2020) |
| mcad | medium chain acyl CoA dehydrogenase | AGAGGACTGTGGTGGAATGG | CCTGCTCCTGGCTCAGTTAC | (Chen et al., 2020) |
| lcad | long chain acyl-CoA dehydrogenase | CAGTGGTCCTGGCTTCTCTC | AAACGTCTTCGCTCCTTGAA | (Chen et al., 2020) |

| | | | | |
|---|---|---|---|---|
| acox1 | acyl-CoA oxidase 1, palmitoyl | ATGCCTGGAACAACACTTCC | AGACCCTTCAGCCTCTGTGA | (Chen et al., 2020) |
| acss2 | acyl-CoA synthetase short chain family member 2 | TCGCTGCTTTGGTAGAAGGT | GCTCTTTCGCAAACTGTTCC | (Chen et al., 2020) |
| dgat2 | diacylglycerol O-acyltransferase 2 | CATGGCATCTTGTGTTTTGG | TCGGTTTACTGGGCAGATTC | (Chen et al., 2020) |
| srebp1 | sterol regulatory element-binding protein 1c | ACTCTTCTGGTGTGGCTGCT | GAGCCTTCAGACACGTCCTC | (Chen et al., 2020) |
| ppara | peroxisome proliferator-activated receptor a | CATCTTGCCTTGCAGACATT | CACGCTCACTTTTCATTTCAC | (Chen et al., 2020) |
| pparb | peroxisome proliferator-activated receptor b | GCGTAAGCTAGTCGCAGGTC | TGCACCAGAGAGTCCATGTC | (Chen et al., 2020) |
| pparg | peroxisome proliferator-activated receptor g | GGTTTCATTACGGCGTTCAC | TGGTTCACGTCACTGGAGAA | (Chen et al., 2020) |
| mttp | microsomal triglyceride transfer protein | CTCAGCTGGTGGATGCAGTA | ATCTCTGTGCTGCCGATCTT | (Chen et al., 2020) |
| apoa1 | apolipoprotein a1 | CCAATTTGTTCCAGGCTGAT | CAACTGGGTGGAGATGGTCT | (Chen et al., 2020) |
| apoc1 | apolipoprotein c1 | AAAGGGACAAGCCATCTGTG | TTTGTGAAATGCTGCTCCAG | (Chen et al., 2020) |
| apoba | apolipoprotein ba | TGACCTCAAGCACGTCACTC | GGGGAAAACCAGCACTTGTA | (Chen et al., 2020) |

**Table S2**. Changes in the growth and development parameters of F1 male zebrafish.

| Parameters | Control | 0.1 μg/L | 1.0 μg/L |
|---|---|---|---|
| Body Weight (mg) | 219.2 ±23.8 | 208.3 ±9.3 | 221.9 ±17.7 |
| BSI | 2.17 ±0.56 | 1.62 ±0.62 | 2.02 ±0.78 |
| HSI | 2.08 ±0.88 | 2.12 ±0.42 | 1.76 ±0.55 |
| GSI | 2.11 ±0.79 | 1.57 ±0.51 | 1.80 ±0.87 |

Note: Brain-somatic index (BSI)=brain weight (mg)×100/body weight (mg); hepatosomatic index (HSI)=hepatic weight (mg)×100/body weight (mg); gonadosomatic index (GSI)=gonad weight (mg)×100/body weight (mg). Values are shown as the mean ± SD of three replicates per treatment (*$p$ < 0.05).

**Table S3**. Biochemical alterations in liver, blood and intestine tissues of F0 female zebrafish.

| Tissues | Indices | Control | 0.1 μg/L | 1.0 μg/L | 10 μg/L |
|---|---|---|---|---|---|
| Liver | TC (mmol/g protein) | 1.64 ±0.12 | 0.66 ±0.10* | 0.38 ±0.03* | 0.65 ±0.28* |
| | TG (mmol/g protein) | 3.49 ±0.16 | 3.50 ±0.52 | 2.64 ±0.37 | 2.10 ±0.66* |
| | FFA (mmol/g protein) | 0.36 ±0.00 | 0.26 ±0.07 | 0.25 ±0.12 | 0.20 ±0.02* |
| | TBA (μmol/g protein) | 6.02 ±1.02 | 4.62 ±0.71 | 3.65 ±0.45* | 4.48 ±0.70* |
| Blood | LDL-C (mmol/L) | 24.38 ±1.29 | 21.67 ±3.34 | 18.87 ±1.49* | 21.40 ±2.10* |
| | HDL-C (mmol/L) | 6.16 ±0.53 | 6.16 ±0.38 | 4.74 ±0.25* | 6.28 ±1.06 |
| | TC (mmol/L) | 7.69 ±0.14 | 8.12 ±0.75 | 8.00 ±0.27 | 6.83 ±0.18* |
| | TG (mmol/L) | 1.15 ±0.07 | 1.03 ±0.07 | 0.91 ±0.04* | 0.86 ±0.09* |
| | TBA (μmol/L) | 4.68 ±0.34 | 4.36 ±0.85 | 3.05 ±0.97* | 2.64 ±0.16* |
| Intestine | TBA (μmol/g protein) | 0.62 ±0.06 | 0.61 ±0.03 | 0.37 ±0.12* | 0.35 ±0.06* |

Note: TC, total cholesterol; TG, triglycerides; FFA, free fatty acids; TBA, total bile acids; LDL-C, low-density lipoprotein cholesterol; HDL-C, high-density lipoprotein cholesterol. Values are shown as the mean ± SD of three replicates per treatment (*$p$ < 0.05).

**Table S4**. Gene transcriptional changes involved in hepatic lipid metabolisms of F0 female zebrafish.

| **Target gene** | Control | 0.1 μg/L | 1.0 μg/L | 10 μg/L |
|---|---|---|---|---|
| *lipca* | 1.00 ±0.03 | 0.55 ±0.33 | 0.58 ±0.40 | 0.23 ±0.17* |
| *lpl* | 1.00 ±0.03 | 0.82 ±0.41 | 1.61 ±1.73 | 0.30 ±0.18 |
| *lipea* | 1.00 ±0.02 | 1.73 ±0.70 | 4.81 ±3.35* | 0.75 ±0.16 |
| *cyp7a1* | 1.00 ±0.03 | 1.38 ±0.08 | 2.91 ±0.67* | 0.37 ±0.08 |
| *cyp8b1* | 1.00 ±0.06 | 1.05 ±0.35 | 0.56 ±0.22* | 0.40 ±0.20* |
| *nr1h4* | 1.00 ±0.01 | 0.98 ±0.27 | 0.74 ±0.05 | 0.44 ±0.27* |
| *cyp27a1* | 1.00 ±0.01 | 1.14 ±0.20 | 0.36 ±0.12* | 0.87 ±0.45 |
| *cyp7b1* | 1.00 ±0.03 | 0.68 ±0.12 | 0.56 ±0.38* | 0.51 ±0.20* |
| *hsd3b7* | 1.00 ±0.02 | 1.17 ±0.15 | 0.47 ±0.13* | 0.48 ±0.12* |
| *bsep* | 1.00 ±0.01 | 1.09 ±0.67 | 1.37 ±1.32 | 1.25 ±0.24 |
| *cs* | 1.00 ±0.04 | 1.33 ±0.21* | 0.47 ±0.10* | 0.46 ±0.23* |
| *aclya* | 1.00 ±0.02 | 0.60 ±0.13 | 1.35 ±0.41 | 0.81 ±0.09 |
| *acaca* | 1.00 ±0.01 | 0.57 ±0.26 | 0.53 ±0.32 | 0.37 ±0.31* |
| *fas* | 1.00 ±0.01 | 3.61 ±4.85 | 2.94 ±3.55 | 0.31 ±0.21 |
| *cpt2* | 1.00 ±0.03 | 0.45 ±0.30* | 0.68 ±0.01 | 0.55 ±0.40 |
| *cpt1* | 1.00 ±0.04 | 1.05 ±0.22 | 1.96 ±0.31* | 0.49 ±0.20* |
| *mcad* | 1.00 ±0.02 | 0.80 ±0.29 | 0.75 ±0.22 | 0.38 ±0.24* |
| *lcad* | 1.00 ±0.03 | 0.47 ±0.14* | 0.72 ±0.24 | 0.42 ±0.23* |
| *acox1* | 1.00 ±0.02 | 0.56 ±0.06 | 0.92 ±0.41 | 0.34 ±0.22* |
| *acss2* | 1.00 ±0.02 | 0.68 ±0.34 | 0.96 ±0.29 | 0.48 ±0.35 |
| *dgat2* | 1.00 ±0.03 | 0.92 ±0.46 | 0.53 ±0.36 | 0.19 ±0.10* |
| *srebp1* | 1.00 ±0.02 | 0.26 ±0.15* | 0.27 ±0.16* | 0.36 ±0.19* |
| *ppara* | 1.00 ±0.00 | 1.24 ±0.29 | 1.50 ±0.93 | 0.40 ±0.07 |
| *pparb* | 1.00 ±0.03 | 0.73 ±0.47 | 0.90 ±0.34 | 0.50 ±0.08 |
| *pparg* | 1.00 ±0.04 | 1.30 ±0.14* | 0.42 ±0.15* | 0.32 ±0.18* |
| *mttp* | 1.00 ±0.04 | 1.09 ±0.28 | 0.67 ±0.12* | 0.63 ±0.13* |
| *apoa1* | 1.00 ±0.01 | 0.67 ±0.56 | 0.29 ±0.06* | 0.13 ±0.06* |
| *apoc1* | 1.00 ±0.02 | 0.72 ±0.26 | 0.64 ±0.49 | 0.22 ±0.19* |
| *apoba* | 1.00 ±0.02 | 0.54 ±0.30* | 0.37 ±0.11* | 0.21 ±0.11* |

**Table S5**. Biochemical alterations in liver, blood and intestine tissues of F0 male zebrafish.

| Tissues | Indices | Control | 0.1 μg/L | 1.0 μg/L | 10 μg/L |
|---|---|---|---|---|---|
| Liver | TC (mmol/g protein) | 0.19 ±0.02 | 0.15 ±0.03* | 0.13 ±0.02* | 0.11 ±0.02* |
| | TG (mmol/g protein) | 1.26 ±0.44 | 1.22 ±0.04 | 0.91 ±0.18 | 0.68 ±0.07* |
| | FFA (mmol/g protein) | 0.28 ±0.07 | 0.22 ±0.06 | 0.16 ±0.02* | 0.18 ±0.02* |
| | TBA (μmol/g protein) | 9.45 ±0.89 | 4.22 ±0.81* | 5.05 ±0.89* | 6.76 ±1.03* |
| Blood | LDL-C (mmol/L) | 31.19 ±1.03 | 24.24 ±0.75 | 23.43 ±1.29* | 16.97 ±7.33* |
| | HDL-C (mmol/L) | 10.4 ±0.05 | 8.65 ±0.74* | 5.94 ±0.74* | 6.03 ±0.24* |
| | TC (mmol/L) | 12.29 ±0.50 | 9.19 ±0.18* | 7.94 ±0.23* | 8.05 ±0.16* |
| | TG (mmol/L) | 0.67 ±0.04 | 0.73 ±0.10 | 0.46 ±0.10* | 0.77 ±0.07 |
| | TBA (μmol/L) | 4.86 ±0.69 | 2.86 ±0.24* | 3.18 ±0.89* | 1.45 ±0.80* |
| Intestine | TBA (μmol/g protein) | 0.57 ±0.08 | 0.58 ±0.17 | 0.34 ±0.12* | 0.32 ±0.09* |

Note: TC, total cholesterol; TG, triglycerides; FFA, free fatty acids; TBA, total bile acids; LDL-C, low-density lipoprotein cholesterol; HDL-C, high-density lipoprotein cholesterol. Values are shown as the mean ± SD of three replicates per treatment (*$p < 0.05$).

**Table S6**. Gene transcriptional changes involved in hepatic lipid metabolisms of F0 male zebrafish.

| **Target gene** | Control | 0.1 μg/L | 1.0 μg/L | 10 μg/L |
|---|---|---|---|---|
| *lipca* | 1.00 ±0.01 | 0.52 ±0.36 | 0.64 ±0.19 | 1.08 ±0.84 |
| *lpl* | 1.00 ±0.02 | 0.64 ±0.51 | 0.80 ±0.33 | 1.28 ±0.65 |
| *lipea* | 1.00 ±0.05 | 1.08 ±1.00 | 1.03 ±0.65 | 1.75 ±1.71 |
| *cyp7a1* | 1.00 ±0.02 | 0.35 ±0.27* | 0.20 ±0.07* | 0.53 ±0.27* |
| *cyp8b1* | 1.00 ±0.01 | 0.57 ±0.26* | 0.39 ±0.17* | 0.50 ±0.29* |
| *nr1h4* | 1.00 ±0.02 | 0.52 ±0.49 | 0.31 ±0.12 | 0.97 ±0.87 |
| *cyp27a1* | 1.00 ±0.02 | 0.57 ±0.54 | 0.52 ±0.22 | 0.67 ±0.46 |
| *cyp7b1* | 1.00 ±0.02 | 0.66 ±0.66 | 1.11 ±0.26 | 1.34 ±0.87 |
| *hsd3b7* | 1.00 ±0.00 | 0.74 ±0.69 | 0.79 ±0.37 | 0.72 ±0.27 |
| *bsep* | 1.00 ±0.01 | 0.41 ±0.17 | 0.56 ±0.25 | 1.31 ±1.43 |
| *cs* | 1.00 ±0.02 | 0.48 ±0.04 | 0.42 ±0.29 | 2.97 ±2.04* |
| *aclya* | 1.00 ±0.02 | 0.45 ±0.33 | 0.60 ±0.29 | 1.86 ±0.45* |
| *acaca* | 1.00 ±0.05 | 0.63 ±0.42 | 0.53 ±0.15 | 2.43 ±2.66 |
| *fas* | 1.00 ±0.03 | 2.15 ±2.97 | 1.51 ±1.18 | 2.88 ±2.82 |
| *cpt2* | 1.00 ±0.05 | 0.40 ±0.24* | 0.20 ±0.06* | 0.52 ±0.20* |
| *cpt1* | 1.00 ±0.02 | 0.91 ±0.68 | 1.25 ±0.65 | 1.25 ±0.85 |
| *mcad* | 1.00 ±0.02 | 0.61 ±0.40 | 0.42 ±0.09* | 0.62 ±0.26 |
| *lcad* | 1.00 ±0.05 | 0.56 ±0.46 | 0.40 ±0.07* | 0.49 ±0.06* |

| | | | | |
|---|---|---|---|---|
| *acox1* | 1.00 ±0.01 | 0.47 ±0.33* | 0.36 ±0.19* | 0.49 ±0.07* |
| *acss2* | 1.00 ±0.07 | 0.37 ±0.14* | 0.52 ±0.27 | 0.91 ±0.53 |
| *dgat2* | 1.00 ±0.03 | 0.79 ±0.84 | 0.54 ±0.27 | 1.41 ±1.37 |
| *srebp1* | 1.00 ±0.03 | 1.87 ±2.59 | 1.19 ±0.69 | 4.08 ±4.23 |
| *ppara* | 1.00 ±0.04 | 0.95 ±1.01 | 1.04 ±0.28 | 1.77 ±1.41 |
| *pparb* | 1.00 ±0.06 | 0.73 ±0.53 | 1.24 ±0.58 | 1.49 ±0.95 |
| *pparg* | 1.00 ±0.07 | 0.29 ±0.06* | 0.50 ±0.15* | 0.47 ±0.07* |
| *mttp* | 1.00 ±0.04 | 0.49 ±0.25* | 0.43 ±0.14* | 0.52 ±0.34* |
| *apoa1* | 1.00 ±0.05 | 0.30 ±0.29* | 0.18 ±0.04* | 0.42 ±0.22* |
| *apoc1* | 1.00 ±0.04 | 0.40 ±0.44 | 0.40 ±0.19 | 0.96 ±0.67 |
| *apoba* | 1.00 ±0.01 | 0.33 ±0.31* | 0.30 ±0.07* | 0.41 ±0.31* |

**Table S7**. Biochemical alterations in liver, blood and intestine tissues of F1 male zebrafish.

| Tissues | Indices | Control | 0.1 μg/L | 1.0 μg/L |
|---|---|---|---|---|
| Liver | TC (mmol/g protein) | 0.59 ±0.17 | 0.62 ±0.14 | 0.88 ±0.20 |
| | TG (mmol/g protein) | 1.67 ±0.49 | 1.42 ±0.42 | 1.67 ±0.17 |
| | FFA (mmol/g protein) | 0.58 ±0.05 | 0.58 ±0.08 | 0.56 ±0.07 |
| | TBA (μmol/g protein) | 5.10 ±1.52 | 5.35 ±3.32 | 11.45 ±2.15* |
| Blood | LDL-C (mmol/L) | 26.08 ±2.31 | 26.22 ±1.71 | 25.95 ±2.57 |
| | HDL-C (mmol/L) | 5.77 ±0.27 | 5.52 ±0.43 | 5.36 ±0.80 |
| | TC (mmol/L) | 13.42 ±1.11 | 13.83 ±1.34 | 13.39 ±0.31 |
| | TG (mmol/L) | 1.96 ±0.05 | 1.46 ±0.07* | 1.72 ±0.09* |
| | TBA (μmol/L) | 3.75 ±0.47 | 5.94 ±1.77 | 6.56 ±0.94* |
| Intestine | TBA (μmol/g protein) | 0.63 ±0.04 | 0.76 ±0.15 | 0.83 ±0.04* |

Note: TC, total cholesterol; TG, triglycerides; FFA, free fatty acids; TBA, total bile acids; LDL-C, low-density lipoprotein cholesterol; HDL-C, high-density lipoprotein cholesterol. Values are shown as the mean ± SD of three replicates per treatment (*$p < 0.05$).

**Table S8**. Gene transcriptional changes involved in hepatic lipid metabolisms of F1 male zebrafish.

| **Target gene** | Control | 0.1 μg/L | 1.0 μg/L |
|---|---|---|---|
| *lipca* | 1.00 ±0.06 | 3.75 ±0.96* | 2.64 ±1.22 |

| | | | |
|---|---|---|---|
| *lpl* | 1.00 ±0.05 | 3.65 ±0.51 | 3.09 ±2.33 |
| *lipea* | 1.00 ±0.02 | 5.85 ±2.78* | 2.05 ±0.84 |
| *cyp7a1* | 1.00 ±0.08 | 13.17 ±10.58* | 0.73 ±0.33 |
| *cyp8b1* | 1.00 ±0.07 | 6.97 ±3.70* | 3.00 ±1.70 |
| *nr1h4* | 1.00 ±0.01 | 4.61 ±2.41* | 2.01 ±0.93 |
| *cyp27a1* | 1.00 ±0.05 | 3.16 ±0.44* | 1.62 ±1.27 |
| *cyp7b1* | 1.00 ±0.08 | 3.60 ±2.04 | 2.77 ±1.43 |
| *hsd3b7* | 1.00 ±0.10 | 1.63 ±0.88 | 1.59 ±1.28 |
| *bsep* | 1.00 ±0.05 | 4.22 ±0.52* | 2.50 ±1.33 |
| *cs* | 1.00 ±0.05 | 4.59 ±3.82 | 1.25 ±1.01 |
| *aclya* | 1.00 ±0.02 | 9.89 ±6.67* | 5.07 ±0.92 |
| *acaca* | 1.00 ±0.03 | 10.58 ±5.79* | 0.85 ±0.50 |
| *fas* | 1.00 ±0.05 | 7.33 ±3.99* | 1.11 ±0.05 |
| *cpt2* | 1.00 ±0.06 | 3.86 ±0.87* | 2.46 ±1.04 |
| *cpt1* | 1.00 ±0.05 | 5.64 ±3.24* | 3.93 ±1.82 |
| *mcad* | 1.00 ±0.05 | 3.55 ±2.33 | 2.03 ±1.23 |
| *lcad* | 1.00 ±0.05 | 3.92 ±1.88 | 3.95 ±3.58 |
| *acox1* | 1.00 ±0.07 | 2.65 ±1.44 | 1.26 ±0.92 |
| *acss2* | 1.00 ±0.01 | 2.49 ±1.13* | 2.24 ±0.39 |
| *dgat2* | 1.00 ±0.05 | 4.74 ±2.50* | 2.72 ±1.94 |
| *srebp1* | 1.00 ±0.09 | 9.48 ±4.11* | 3.31 ±0.65 |
| *ppara* | 1.00 ±0.08 | 2.93 ±2.16 | 1.59 ±0.73 |
| *pparb* | 1.00 ±0.08 | 5.49 ±0.62* | 3.89 ±1.51* |
| *pparg* | 1.00 ±0.08 | 3.54 ±1.06* | 2.01 ±1.28 |
| *mttp* | 1.00 ±0.02 | 2.72 ±1.05* | 1.82 ±0.87 |
| *apoa1* | 1.00 ±0.02 | 10.25 ±6.97 | 8.37 ±4.55 |
| *apoc1* | 1.00 ±0.08 | 5.68 ±0.41* | 4.46 ±3.53 |
| *apoba* | 1.00 ±0.06 | 6.84 ±1.64* | 3.55 ±2.32 |

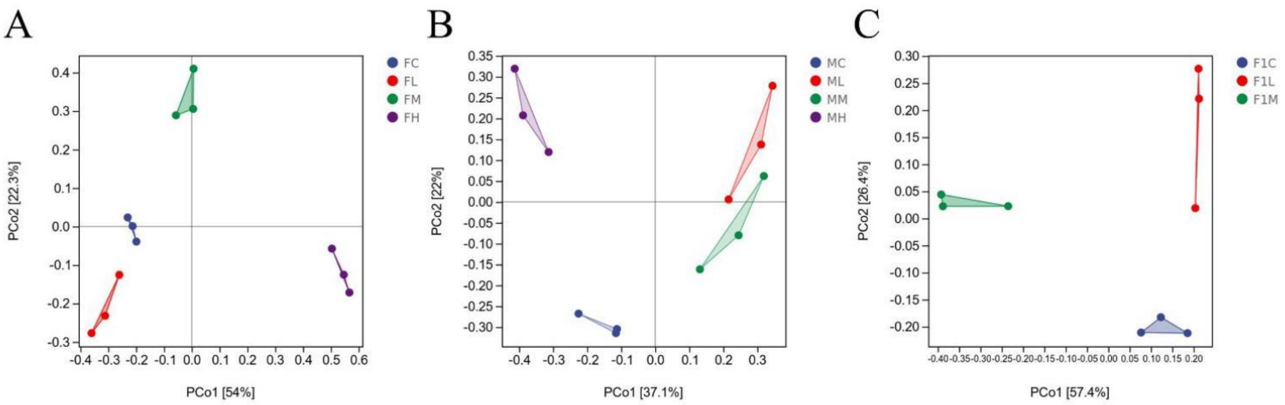


**Figure S1**. Principal coordinate analysis (PCoA) of the gut microbiome of F0 and F1 generations. A: Gut microbiome of F0 female fish (FC: female fish in control group; FL: female fish in 0.1 μg/L DCZ group; FM: female fish in 1.0 μg/L DCZ group; FH: female fish in 10 μg/L DCZ group) ;B: Gut microbiome of F0 male fish (MC: male fish in control group; ML: male fish in 0.1 μg/L DCZ group; MM: male fish in 1.0 μg/L DCZ group; MH: male fish in 10 μg/L DCZ group); C: Gut microbiome of F1 male fish (F1C: male fish in control group; F1L: male fish in 0.1 μg/L DCZ group; F1M: male fish in 1.0 μg/L DCZ group).

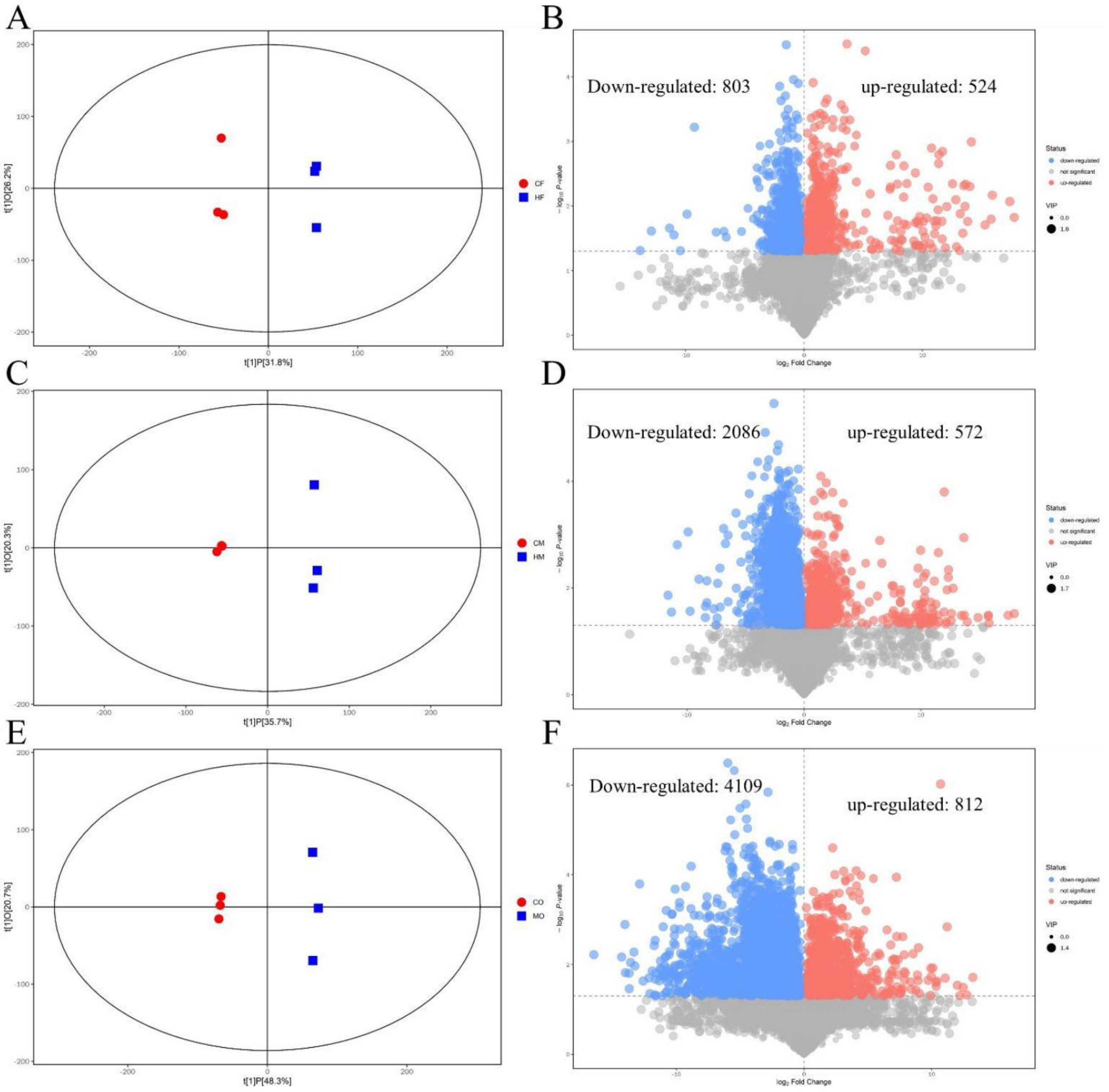


**Figure S2**. OPLS-DA plot and volcano plot of DCZ exposure on intestinal metabolism of F0 and F1 generations. A, C and E are the OPLS-DA analysis of the gut differential metabolites of F0 generation female fish, F0 generation male fish and F1 generation male fish, respectively; B, D and F are volcano plots of the gut differential metabolites of F0 generation female fish, F0 generation male fish and F1 generation male fish, respectively. CF: F0 female fish-control; HF: F0 female fish-10 μg/L DCZ; CM: F0 male fish-control; HM: F0 male fish-10 μg/L DCZ. CO: F1 male fish-control; MO: F1 male fish-1.0 μg/L DCZ.

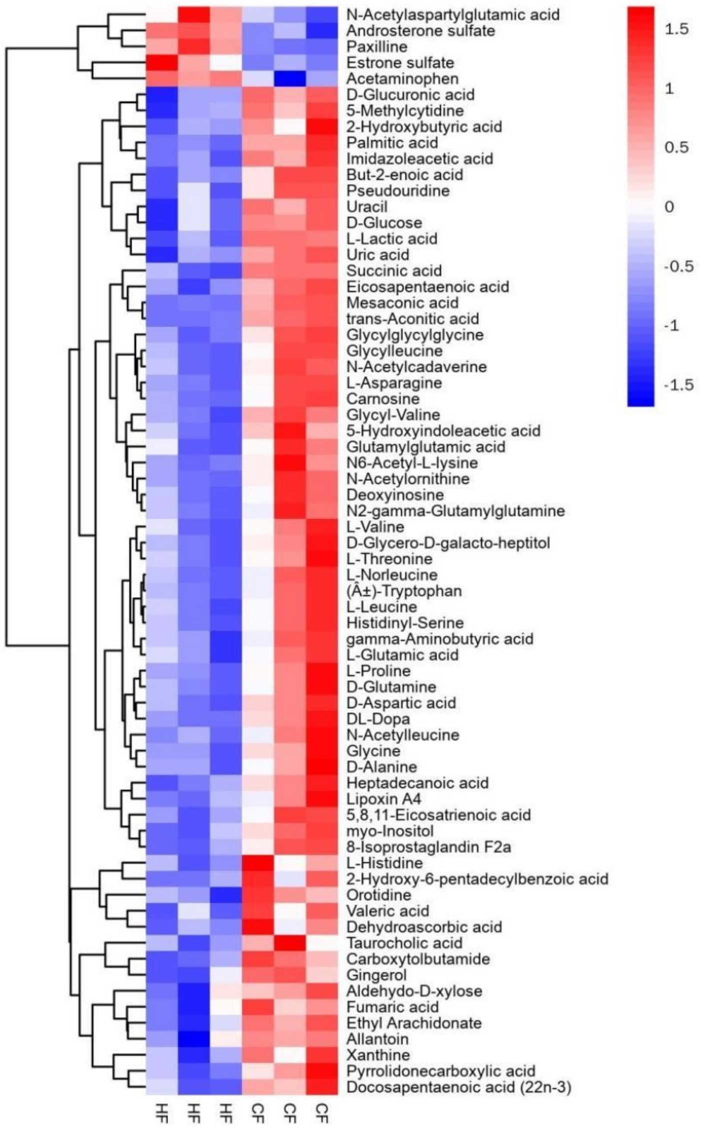


**Figure S3**. Heat map of differential metabolites in F0 female fish gut.

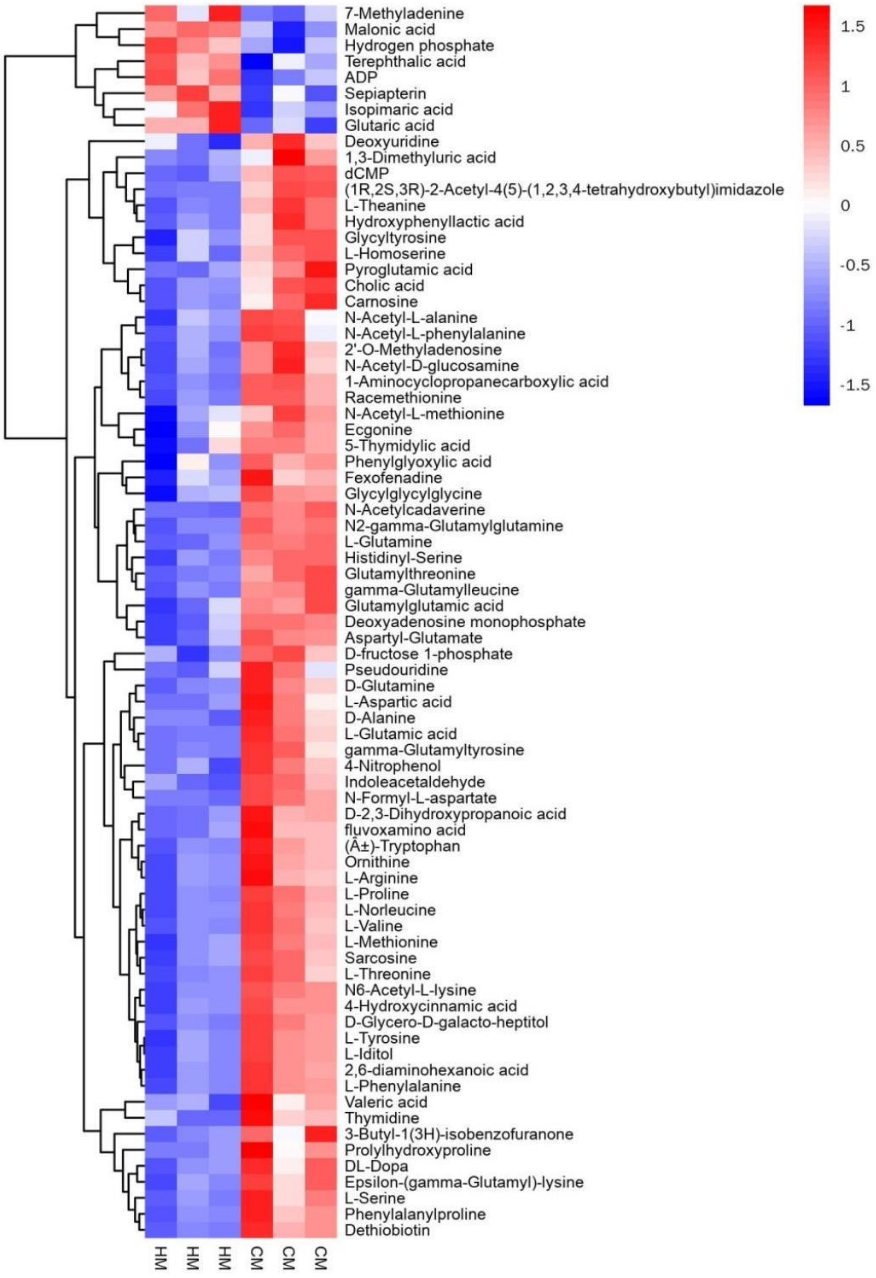


**Figure S4**. Heat map of differential metabolites in F0 male fish gut.

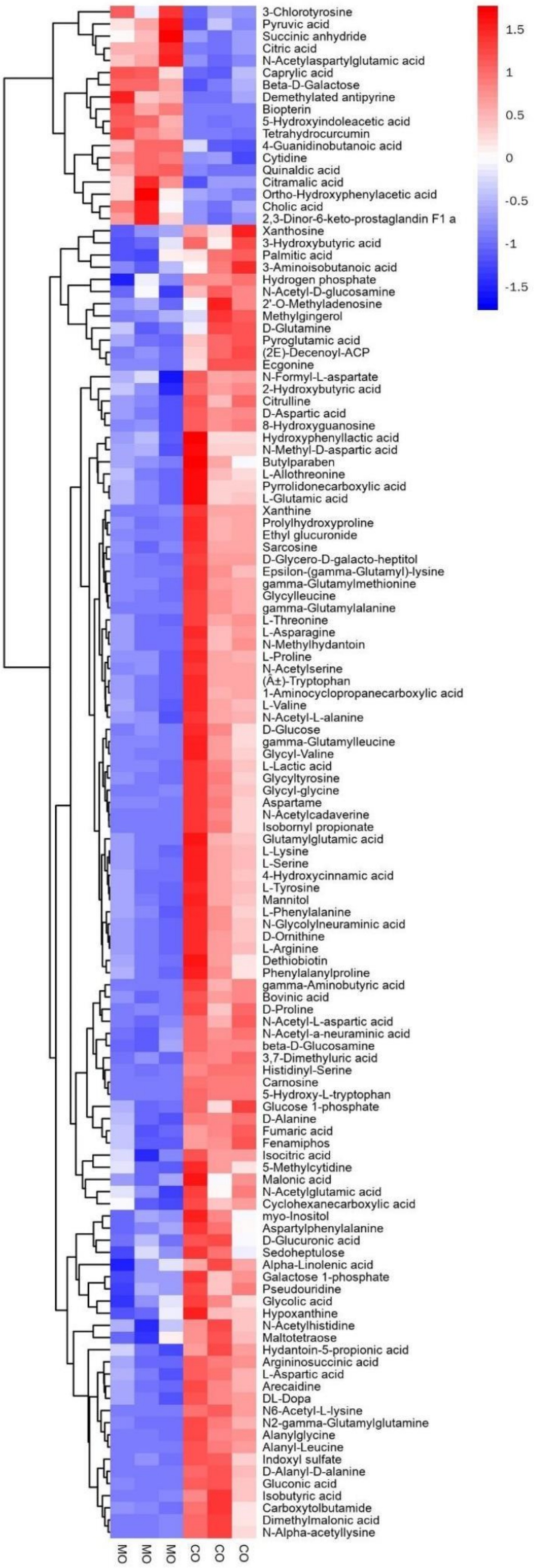


**Figure S5**. Heat map of differential metabolites in F1 male fish gut.
